# Visual Comparison of Phylogenetic Trees Through iPhyloC, a New Interactive Web-Based Framework

**DOI:** 10.1101/2021.05.14.444083

**Authors:** Muhsen Hammoud, Charles Morphy D. Santos, João Paulo Gois

## Abstract

Current side-by-side phylogenetic trees comparison frameworks face two issues: (1) accepting binary trees as input, and (2) assuming input trees having identical or highly overlapping taxa. We present a task abstraction of the problem of side-by-side comparison of two phylogenetic trees and propose a set-based measure for detailed structural comparison between two phylogenetic trees, which can be non-binary and not highly overlapping. iPhyloC is an interactive web-based framework including automatic identification of the common taxa in both trees, comparing input trees in several modes, intuitive design, high usability, scalability to large trees, and cross-platform support. iPhyloC was tested in hypothetical and real biological examples.

## 1. Introduction

Evolution produces the natural hierarchy among species, groups of species and genes that can be represented through rooted or unrooted dendograms ^3^. There are tools for phylogenetic tree inference such as TNT ^9^ and MESQUITE ^16^, where the analysis of a single-source data set may result in dozens, or even hundreds, of equally most parsimonious trees, *i*.*e*., trees with the same minimum number of steps. All of these trees compose the so-called *tree space*.

Comparing trees derived from different data-sets is extremely useful. However, the taxon sampling can be biased depending on the sort of primary evidence used in the phylogenetic analysis, which may lead to trees seeming incomparable at first. There are some molecular-based phylogenies with scarce representation of uncommon or hardly sequenced taxa (as fossil species) ^8^. Comparing such molecular based-trees with a morphological-based tree in search for common natural groups and stable phylogenetic relationships is not straightforward.

We identify two main limitations for biological systematics of the current phylogenetic trees visual comparison frameworks: (1) accepting binary trees as input; and (2) assuming input trees having identical, or at least highly overlapping, sets of taxa. Such assumptions prevent biologists from using these frameworks to compare phylogenetic trees that do not fulfill these limitations, which is the case, for instance, in the comparison of a phylogenetic supertree with its source trees (often the supertree is not totally resolved and do not highly overlap with its source trees). We address the aforementioned restrictions by introducing a phylogenetic trees comparison framework which accepts binary and non-binary trees as input, regardless of their overlapping level. This work provides (1) task and data abstraction for trees comparison; (2) an interactive visual comparison framework to compare two trees side-by-side named iPhyloC; and (3) validation through usage scenarios.

## 2. Related work

Liu et al. ^15^ divided the trees visual comparison frameworks in *Few in Full, Dozens at Multi-Scale*, and *Many as Points*. iPhyloC belongs to the *Few in Full* category, which comprises systems that handle small number of trees (often two), making them very scalable since they can deal with a massive number of nodes per tree. Specifically, iPhyloC handles two phylogenetic trees at the same time. The main difference between ours and the current available frameworks is the ability to compare non-binary and non-highly overlapping phylogenetic trees.

Several systems and packages deal with trees comparison ^20,10^. Phylo.io ^23^ is a web application to visualize and compare two phylogenetic trees side-by-side. Beck et al. ^4^ utilizes superposition to stack trees visually. In addition, two packages for the R programming language - phytools ^21^ and ggtree ^28^ - allow visual comparison of trees. Phytools uses the cophylo function, while ggtree offers plotting functionality of several phylogenetic trees in the same space to facilitate comparison, along with annotation functionality. Although useful, both packages have limitations. The cophylo function in Phytools only matches the tips of the two input trees whereas the ggtree is a tree visualization package; to use it for tree comparison, the user has to plot the trees using R programming, then connect the common taxa through line drawing commands. Both packages do not provide any type of interactive exploration. Finally, they require R programming language and understanding its syntax, which might be time consuming and out of the scope for some biologists.

## 3. Phylogenetic tree data

A phylogenetic tree is a dendogram composed of hierarchically structured set of leaf nodes, the *taxa*. The internal nodes of a tree represent common ancestors. A *clade*, or a *monophyletic group*, is the set of all taxa underneath a specific internal node including the common ancestor, while a *subtree* is the set of all descendants beneath an ancestor including the hierarchical structure, *i*.*e*., the internal nodes. In biological research, systematists usually compare (1) a reference tree with a collection of other trees to test the main hypothesis, (2) a collection of trees without having a reference tree, or (3) trees side-by-side, which is the case of iPhyloC.

As input, iPhyloC accepts two phylogenetic trees, 𝒯_1_ and 𝒯_2_, in parenthetical format, with or without branch length, where leaf nodes correspond to taxa and have names, and inner nodes are not labeled. The two input trees can be totally resolved (binary or dichotomous trees) or partially resolved (non-binary trees, where polytomies are present). It is not necessary for the terminal taxa to be the same in both trees (non-highly overlapping sets are particularly useful since most part of the available tools for tree comparison deal only with trees composed by the same set of taxa). 𝒯_1_ and 𝒯_2_ should not have paralog terminals. Figure 1 depicts the binary tree, non-binary tree, and tree with paralogs concepts.

**Fig. 1:**
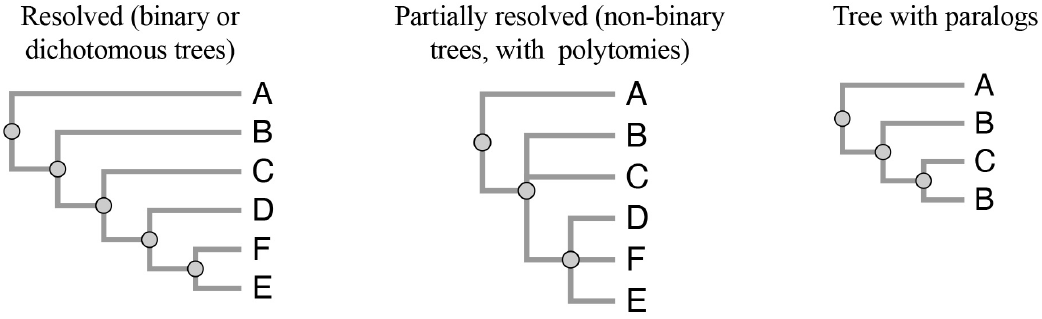
Types of phylogenetic trees.

## 4. Task abstraction

The goal of iPhyloC is to compare two phylogenetic trees side-by-side in search for common elements and structural differences. We consider multi-strand approach to task generation ^13^, which means deriving tasks based on several sources including a primary source, interviews with domain experts (a co-author of this paper is a biologist), and a secondary source, from literature ^17,23,25,15^. iPhyloC deals with:

1. **Easily discover the non-shared taxa between both trees:** before starting an in-depth comparison between the two input trees 𝒯_1_ and 𝒯_2_, it is often necessary to identify their degree of similarity, as well as to see the distribution of the shared taxa set 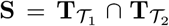, where **T** denotes the taxa set of the phylogenetic trees 𝒯_*i*_, *i* = 1, 2. Giving such a preliminary view on 𝒯_1_ and 𝒯_2_ saves time and effort because it helps the user to decide whether to continue with in-depth comparison or not and on which parts of 𝒯_1_ and 𝒯_2_ to focus.
2. **Compare two trees in multiple modes:** presenting 𝒯_1_ and 𝒯_2_ in several modes eases the comparison process. For example, pruning the non-shared taxa from 𝒯_1_ and 𝒯_2_ helps focusing on the shared taxa **S** by eliminating the noise that non-shared taxa causes. We define tree modes for 𝒯_1_ and 𝒯_2_ as follows: original input tree, non-shared taxa collapsed, and non-shared taxa pruned. Each mode has two states: original taxa order, and alphabetical order. An additional mode is comparing each of the six aforementioned options (three modes with two states for each mode) with the strict consensus tree derived from 𝒯_1_ and 𝒯_2_ after pruning the non-shared taxa, where strict consensus tree summarizes all of the information contained in a set of trees whose taxa are all the same.
3. **Explore the corresponding subtree:** this task means that the user can select any node (internal or leaf node) from one tree (𝒯_1_ or 𝒯_2_) and explore the corresponding subtree in the other tree. The corresponding subtree exploration should be available for all of the comparison modes mentioned in task 2.
4. **Explore each tree separately:** provides the ability to separately interact with each of the trees. The possibilities of exploring the trees include: changing the layout between linear and radial; manipulating the tree layout for better visualization; showing and hiding specific nodes or subtrees; selecting a node, a branch, or a subtree; and re-rooting the tree at a specific node.
5. **Annotate the phylogenetic trees:** provides the user with a set of shapes and text elements that allows the addition of annotations to both trees.

## 5. iPhyloC

To present a preliminary view of 𝒯_1_ and 𝒯_2_, iPhyloC starts with three pre-processing steps, which are: trees pruning, finding the strict consensus tree for the two input trees, and taxa alphabetical ordering. After finishing the three steps, iPhyloC shows an automatic preliminary comparison between 𝒯_1_ and 𝒯_2_.

### 5.1. Phylogenetic trees pre-processing

1. **Trees pruning:** All of the non-shared taxa between the two trees subject to comparison are permanently removed. First, we find the shared taxa **S**. Then, for each tree 𝒯_*i*_, we prune the non-shared taxa **S**^**C**^. The resultant trees are 𝒯_1*P*_ and 𝒯_2*P*_. The pruning process includes removing all inner tree nodes with only one child. Figure 2 exemplifies the tree pruning process.
2. **Finding strict consensus among the input trees:** Consensus methods differ depending on the context in which they are used. When dealing with multiple trees, strict consensus constructs a tree containing only the components shared by all trees ^7^. We focus on the strict consensus method ^12^ because the main goal of our framework is to allow the user to emphasize the congruent, *i*.*e*., evolutionary meaningful, phylogenetic relationships. Computing strict consensus between trees derived from different datasets and taxon sampling – and not among equally most parsimonious trees resultant from a single analysis – is not an option for most of the available phylogenetic software. In our approach, we find the strict consensus tree of 𝒯_1*P*_ and 𝒯_2*P*_ which is 𝒯_*C*_ to provide an overall estimate of the pruned trees. Figure 3 depicts the strict consensus process.
3. **Trees taxa alphabetical ordering:** one of the difficulties of phylogenetic trees comparison is finding the shared terminal taxa in different cladograms, especially the large ones. To facilitate the comparison, we show 𝒯_1_ and 𝒯_2_ with taxa alphabetically ordered as much as possible without changing the internal rela-tionships, while avoiding any edge crossings. This process eases the comparison by helping the user to focus on the actual biological similarities and differences between the two input trees. This approach works only if the branch lengths are identical (if the trees have branch lengths). We provide taxa alphabetical or-der for: 𝒯_1_, 𝒯_1*P*_, 𝒯_2_, 𝒯_2*P*_, and 𝒯_*C*_ named respectively: 𝒯_1*O*_, 𝒯_1*P O*_, 𝒯_2*O*_, 𝒯_2*P O*_, and 𝒯_*CO*_. Figure 4 depicts the taxa ordering process.
4. The results of the pre-processing steps are four variations of each tree: the original uploaded tree (𝒯_*i*_), the original tree with taxa ordered alphabetically (𝒯_*iO*_), a tree containing only the shared taxa with the other tree in the original taxa order (𝒯_*iP*_), and a pruned tree with alphabetically ordered taxa (𝒯_*iP O*_), where *i* ∈{1, 2}; along with that, the strict consensus tree of the pruned trees considering the original taxa order (𝒯_*C*_) and the alphabetical taxa order (𝒯_*CO*_).

### 5.2. Structural comparison

We present here the available structural comparison facilities that iPhyloC offers to the user after the pre-processing step.

**Fig. 2:**
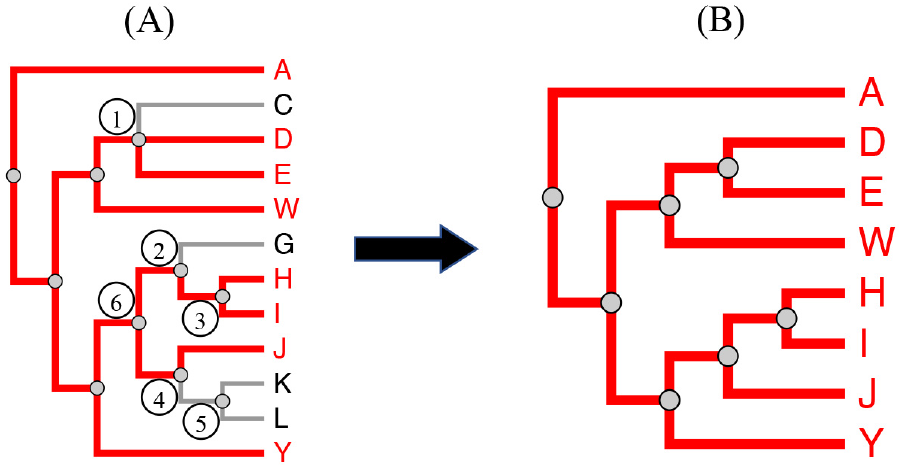
Phylogenetic trees pruning. From tree **(A)**, the taxa **C, G, K**, and **L** undergo the pruning process. When pruning **C**, the inner node **1** is not affected. However, when pruning **G**, the inner node **2** is also pruned because it has only the inner node **3** left. The same happens when pruning **K** and **L**: node **5** is pruned because it has no more child nodes; hence, inner node **4**, with only one child left, **J**, is also pruned. As result **(B)**, inner node **6** becomes directly related to **J**.

**Fig. 3:**
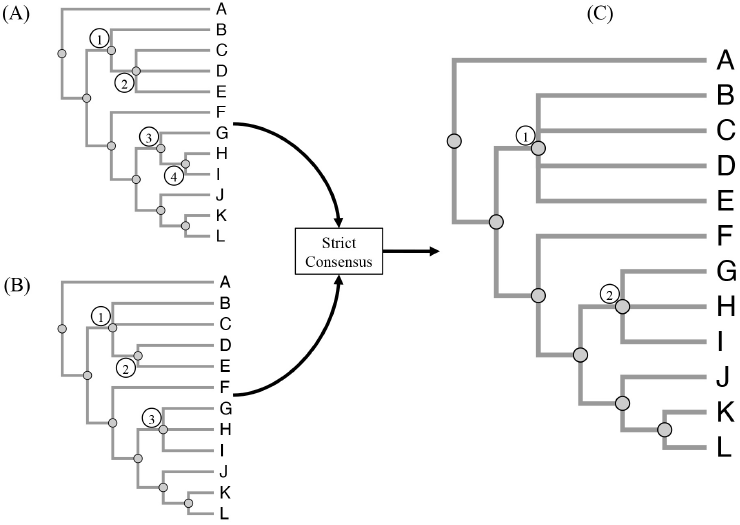
Strict consensus. Trees **(A)** and **(B)** differ in topology but have the same taxa. The consensus tree **(C)** contains only the groups occurring in both **(A)** and **(B)**. Inner node **1** in tree **(C)** is a summarization of inner nodes **1** in tree **(A), 2** in tree **(A), 1** in tree **(B)**, and **2** in tree **(B)**. Similarly, inner node **2** in tree **(C)** is a summarization of inner nodes **3** in tree **(A), 4** in tree **(A)**, and **3** in tree **(B)**.

**Fig. 4:**
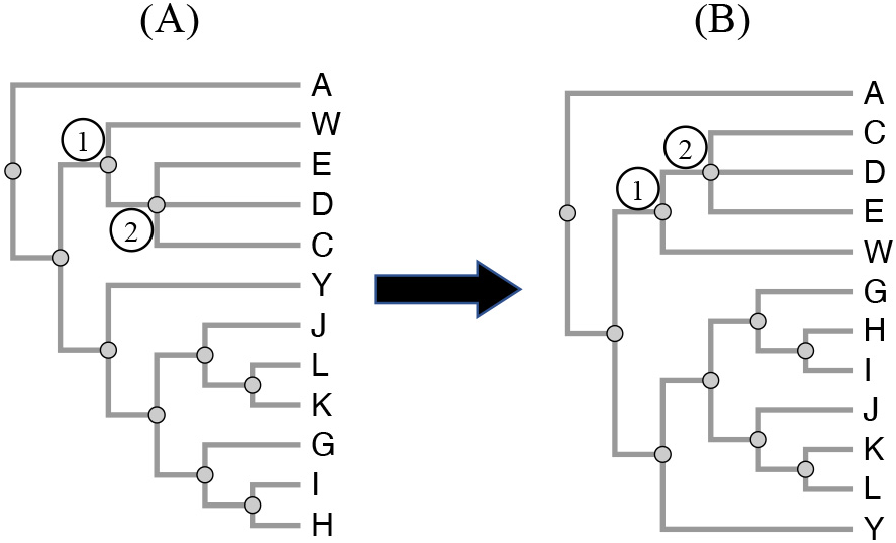
Phylogenetic tree ordering. The trees **(A)** and **(B)** contain the same information. Tree **(B)** is the result of ordering the taxa of tree **(A)**.

1. **Shared tree highlighting:** using 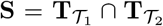, we highlight in 𝒯_1_ and 𝒯_2_ the terminal nodes *t* ∈**S**, and all their related ancestors up to the root node. Figure 5(A) exemplifies shared tree highlighting.
2. **Collapsing non-shared taxa:** unlike the pruning process, collapsing nodes means hiding them without permanent removal. We collapse the non-shared taxa set **S**^**c**^ in 𝒯_1_ and 𝒯_2_, as shown in Figure 5(B) and (C).
3. **The Corresponding SubTree (CST):** finding comparable counterparts between the trees, namely corresponding subtrees (CST), depends on the user’s request. CST is the most similar subtree in the other tree. The user can select a node in the tree displayed on the left side of the screen (𝒯_1_), then iPhyloC will find the corresponding subtree in the tree displayed on the right side (𝒯_2_), and *vice versa*. Figure 6 exemplifies the CST.

CST is a set-based measure for real-time interaction. Unlike similarity metrics that consider the topological differences, such as Robinson-Foulds distance metrics ^22,5^, we consider only the set of leaf nodes from an ancestor (corresponding to a *clade* or a *monophyletic group*).

**Fig. 5:**
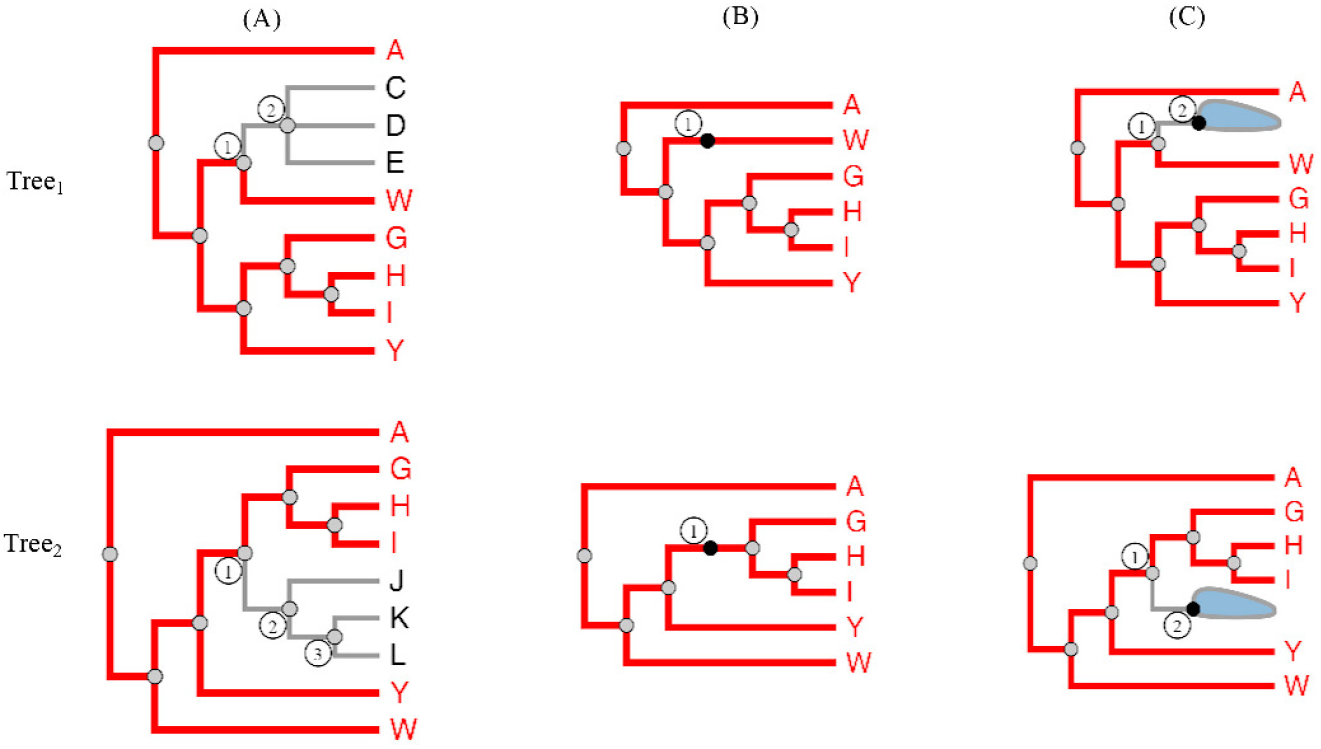
Structural comparison. (A) Highlighting the shared taxa between **Tree**_**1**_ and **Tree**_**2**_ (red color). Taxa **C, D**, and **E** are present in **Tree**_**1**_ only; their ancestor, inner node 2 in **Tree**_**1**_, is not highlighted. In **Tree**_**2**_ taxa **J, K**, and **L** are exclusive; inner nodes 2 and 3 in **Tree**_**2**_ are not highlighted. (B) Non-shared taxa collapsed. The black colored points at inner node 1 in **Tree**_**1**_ and **Tree**_**2**_ indicate the location of hidden nodes. (C) Shows a hidden inner node with a set of leaf nodes.

**Fig. 6:**
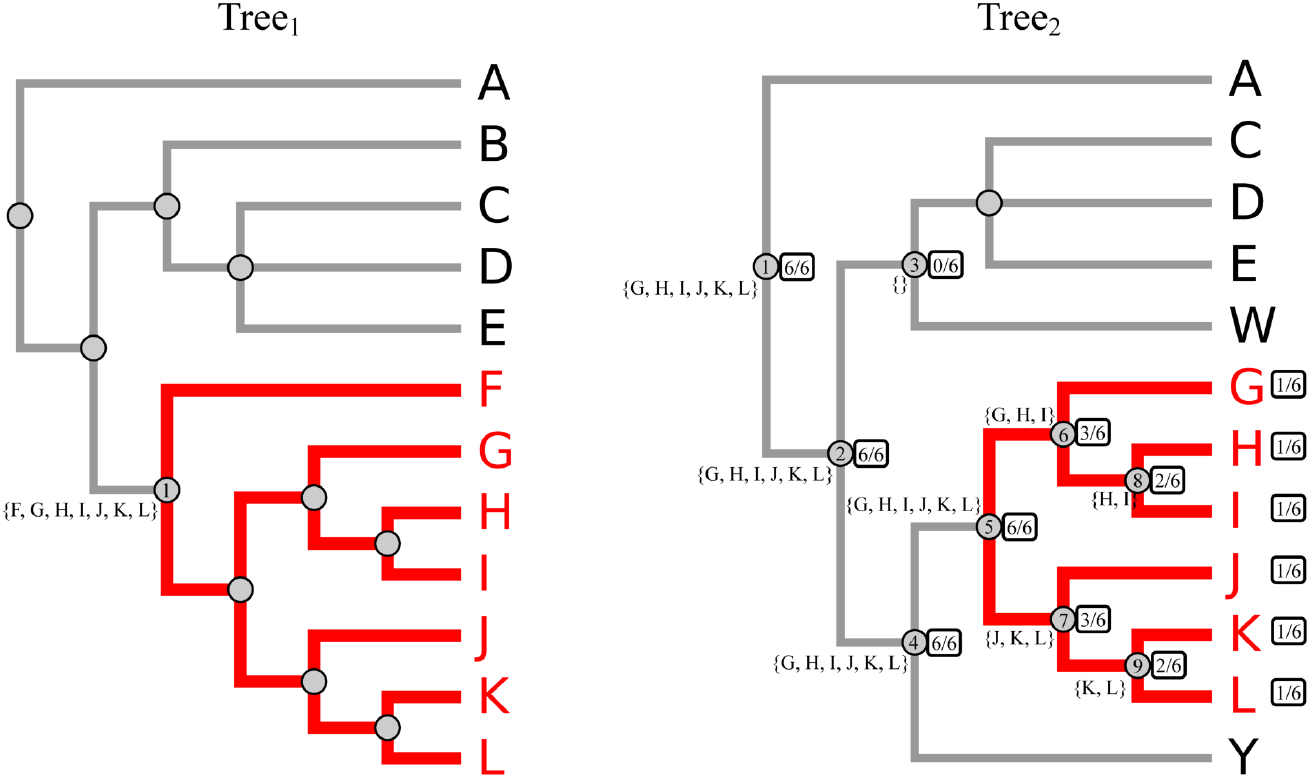
Finding the corresponding subtree (CST). Details in the text.

To facilitate the calculations of CST, we find and store in advance the set of taxa 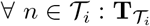 where *i* ∈*{*1, 2}and *n* denotes an inner node of 𝒯_*i*_. Let *n*_1_denote the inner node that the user selected from 𝒯_1_ and 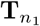 denote the clade of *n*_1_, as in Figure 6. Then the set of shared nodes between the subtree rooted at *n*_1_ and 𝒯_2_ is 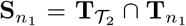. We use 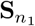 along with breadth first search to find CST in 𝒯_2_. The similarity index is computed as:

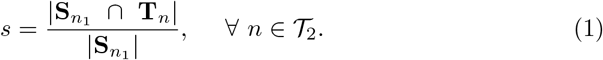

The search finishes in a specific subtree from 𝒯_2_ when *s* = 0. The last step discards all of the nodes *n* ∈𝒯_2_ where *s* = 1 except for the one with the highest depth calculated from the root node of 𝒯_2_.

The second case of CST is when the user chooses a taxon, denoted as **t**. In the second case, iPhyloC will not calculate the *s* index, it will only search if the selected taxon from 𝒯_1_ exists in 𝒯_2_. If **t** exists in 𝒯_2_, then it will be highlighted.

When the user selects node **1** in 𝒯_1_, iPhyloC will search for the CST in 𝒯_2_. *s* is calculated according to equation 1 as shown at the right side of each node in 𝒯_2_. The search stops in a specific subtree if *s* = 0 as in node **3** in 𝒯_2_. We keep only one node with *s* = 1, the one with the maximum depth from the root of the tree, node **5** in 𝒯_2_, and discard other with *s* = 1. Node **5** in 𝒯_2_ represents the root of the corresponding subtree of node **1** in 𝒯_1_.

### 5.3. Trees visualization

Figure 7 shows both linear and the radial layouts in iPhyloC. We kept the design of CST as simple and intuitive as possible to facilitate the exploration process rather than cluttering the visualization with too much information. Each node in the CST has a similarity index *s* > 0, and we use size and color as visual encoding for *s*. Each node in the phylogenetic tree is visualized as a circle with *3px* radius. The nodes of CST have a similarity index *s* ∈[0, 1]. The nodes sizes are normalized using the following equation:

**Fig. 7:**
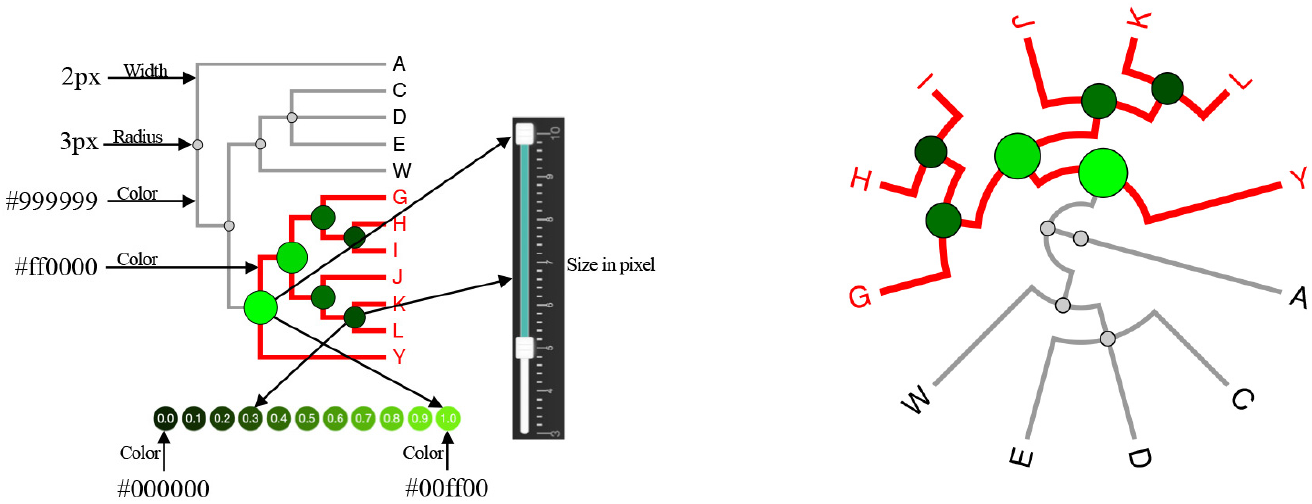
Linear tree layout (left side) and radial tree layout (right side) visualization. The user can control the radius of the CST nodes using a double handles slider. The color scale of the CST nodes is shown at the bottom.

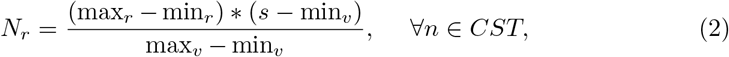

where *N*_*r*_ refers to the node’s radius in pixels, and max_*r*_ and min_*r*_ refer to the maximum and minimum radii. The default values are min_*r*_ = 5 and max_*r*_ = 10, but the user can change these values interactively using a double handles slider as shown in Figure 7. Both max_*v*_ and min_*v*_ refer to the maximum and minimum value of the similarity index, in our case *s* ∈[0, 1]. CST nodes fill color is normalized similarly between the black color and the green color corresponding to similarity values *s* = 0 and *s* = 1 respectively. The user can choose to encode the similarity index values using the aforementioned color scale, or using the default color for all nodes, which is gray.

### 5.4. Trees annotation

Our design choice for annotations is to provide an easy-to-use tool, allowing the user to edit shapes, namely rectangle and arc, along with a text element. iPhyloC’s annotation functionality is not present in any other phylogenetic trees comparison framework currently available. It is based on the use of a rectangle, an arc, and text elements. Figure 8 shows the available shapes and how to edit them.

**Fig. 8:**
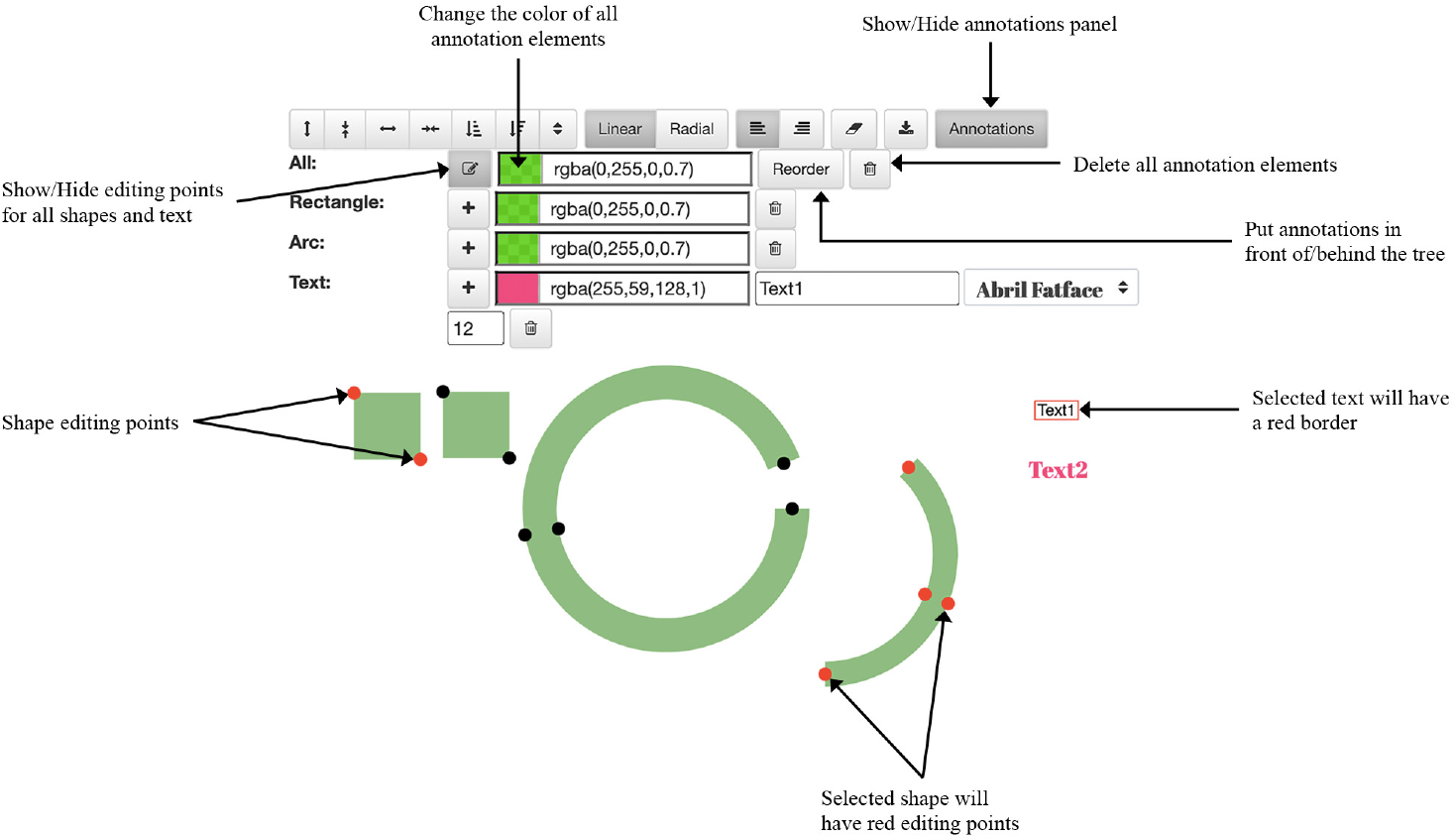
Annotations functionality in iPhyloC

A new shape (a rectangle or an arc) or text element is added in the top-left corner and moved to the tree visualization area. Changing a shape size is done by dragging the small red circles attached to it (the shape’s editing points). Additionally, font type, size, color, and contents are editable. Deleting a shape or a text element is done by selecting it (clicking on it), and then clicking on the delete button in the tools bar. Furthermore, the user can change the color of all shapes and text elements, delete them, and put them in-front-of/behind the phylogenetic tree using the buttons available in the annotations tool bar. The “Reorder” functionality is important because the SVG visualizes its elements in layers, and the user can interact only with the top layer. Consequently, to allow the user to interact with the phylogenetic tree while it is annotated, and to avoid the annotation shapes to block or blur the tree, we have to change the order of the SVG layers.

## 6. Results

We deployed iPhyloC on cloud server running Ubuntu 18.04.5 with 2 Intel Xeon processors, and 7.6 GB memory. iPhyloC can be accessed through the following link http://nuvem.ufabc.edu.br/iphyloc/.

Here, we compare iPhyloC to Phylo.io ^23^ in an usage scenario that is only superficially discussed in the literature, although very important: the phylogenetic trees comparison in the context of supertrees. A supertree is a unique, usually large, phylogenetic tree assembled from a combination of smaller phylogenetic trees, which may have been based on different datasets or different taxa sampling ^1^.

First, we constructed a phylogenetic supertree based on three source trees with different numbers of terminal nodes (taxa) ^27,26,14^ using Fitch parsimony analysis ^24^. Through BuM ^11^, we generated the combined MRP-matrix (which refers to the Matrix Representation with Parsimony ^2^ that is used to generate the supertree). The resultant supertree consists of 146 taxa as shown in Figure 9-iPhyloC 𝒯_2_. The importance of testing this scenario is related to the very nature of a phylogenetic supertree. As aforementioned, sometimes there are little overlap between the supertree and their source trees; moreover, not all of the internal nodes of a supertree are totally resolved, and polytomies are common. Current phylogenetic trees comparison frameworks do not consider this case. For the remainder of this usage scenario, we compare the tree from Ševčík et al. ^26^ (𝒯_1_) with the supertree (𝒯_2_).

**Fig. 9:**
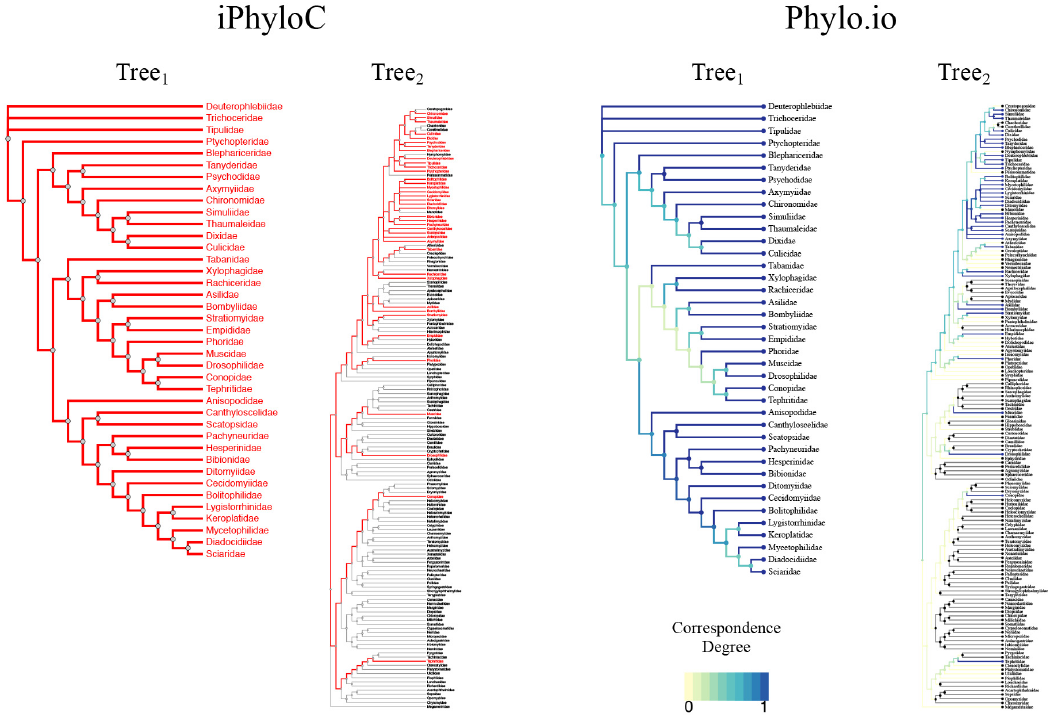
A comparison between iPhyloC and Phylo.io ^23^. We use only two colors in iPhyloC; the dusty gray and red. Phylo.io uses a scale of colors to visualize the correspondence degree

Figure 9 shows the first view of 𝒯_1_ and 𝒯_2_ in both frameworks, the one proposed here, iPhyloC, and Phylo.io ^23^. We use two colors in iPhyloC: dusty gray to visualize the branches of non-shared taxa or their inner nodes, and red to visualize the branches of the shared taxa or their inner nodes. This gives the user a fast and clear idea about the general similarities between 𝒯_1_ and 𝒯_2_. Differently, Phylo.io ^23^ uses a scale of colors starting from yellow to blue. This color scale represents the degree of correspondence of a branch calculated according to the Best Corresponding Node (BCN) index ^17^. Having to interpret several colors when looking at the non-shared taxa between 𝒯_1_ and 𝒯_2_ is not straightforward.

In iPhyloC, the branches of the non-shared taxa are painted gray and easily identified. Phylo.io uses two colors to visualize non-shared taxa (yellow and gray). An example is the last taxon in the bottom of 𝒯_2_, Megamerinidae, is not shared with 𝒯_1_, but the branch that goes back to its direct ancestor is yellow, while the branch of another non-shared taxa (e.g., Chyromyidae) is gray. The task of finding shared and non-shared taxa is unambiguous in iPhyloC, but it is hard in Phylo.io as shown in Figure 9.

Further in-depth comparison reveals the limits of Phylo.io to find the BCN, which is calculated for each node before visualizing 𝒯_1_ and 𝒯_2_, along with a set of interactions to manipulate each tree separately. In iPhyloC, we offer the corresponding sub-tree (CST) instead of BCN. Figure 10 shows an example of CST in iPhyloC where we compare the fully expanded tree from ^26^ with the supertree having all non-shared taxa collapsed. We selected a node from 𝒯_1_, and iPhyloC showed its corresponding subtree in 𝒯_2_ using the node size and the color scale that are shown in the bottom of the figure to encode the correspondence degree of each inner node of the CST. iPhyloC offers the ability to mirror the right-hand tree as well.

**Fig. 10:**
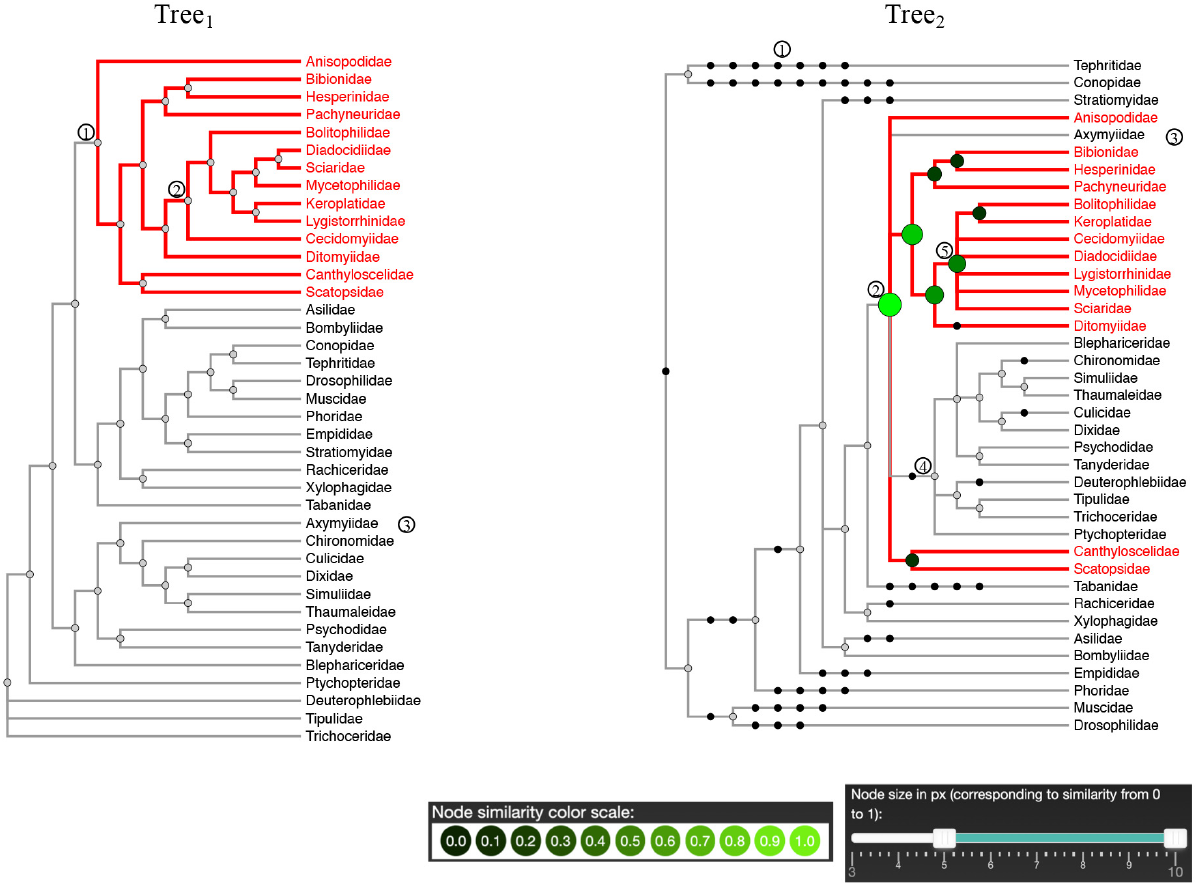
Comparing two trees of Diptera, after collapsing the non-shared leaf nodes, as in node **1** in **Tree**_**1**_, and **Tree**_**2**_, a supertree. This figure shows the corresponding subtree of node **1** in **Tree**_**1**_, which is rooted at node **2** in **Tree**_**2**_. The user can notice the structural differences between node **2** in **Tree**_**1**_ and node **5** in **Tree**_**2**_.

Phylo.io does not provide the radial tree layout, which is especially important when exploring large-scale trees (with 100 or more taxa). On the other hand, Figure 11 shows the radial tree layout in iPhyloC and how it eases the process of highlighting common elements in large trees.

**Fig. 11:**
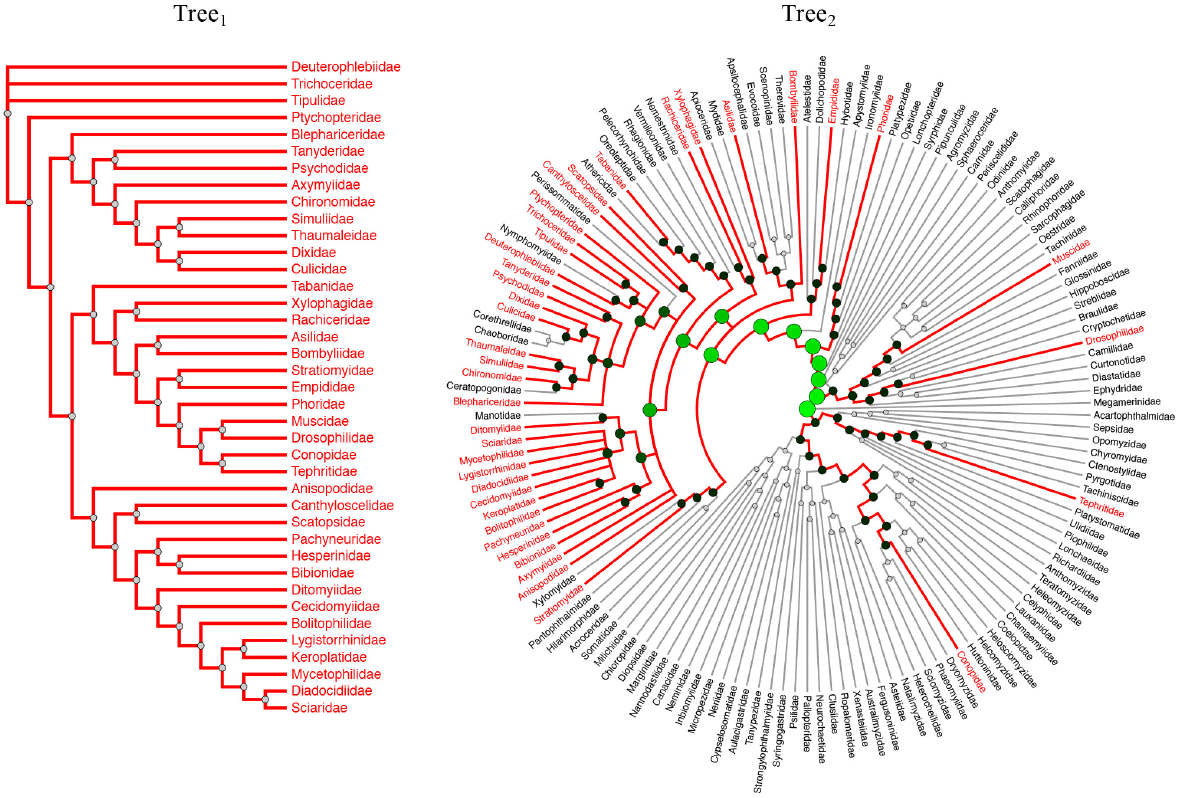
Linear and radial tree layouts. The radial layout is specifically beneficial with large trees (**Tree**_**2**_ in this example consists of 146 taxa).

## 7. Discussion

The main goal of every phylogenetic analysis is to identify monophyletic groups. In this sense, iPhyloC is especially helpful for exploratory analysis, allowing the identification of clades stable enough to be present in several different phylogenies with similar composition of terminals and phylogenetic relationships within them, suggesting that such groups are natural ones and not artifacts of a classification system. The correspondence of phylogenetic patterns among different trees, as visualized by iPhyloC, would help implementing a sort of evaluative “criterion of reality” of a phylogenetic tree as a scientific theory ^6^.

Another interesting issue may raise with iPhyloC comparisons. Even if the relationships of two sets of similar terminals are not correspondent, this may be interpreted as a positive result, since it indicates the need for additional systematics studies for unveiling more robust evolutionary scenarios. Such a feature is also useful for educational purposes, especially for showing the students that the scientific knowledge concerning phylogenetic hypothesis is transient, as any other scientific theory ^6^, and depends on increased amounts of reliable phylogenetic signal.

With the popularization of phylogenetic analysis based on massive amounts of genetic data, software performance is becoming an important issue in biological systematics. We faced the difficulty of addressing the amount of speedup when reviewing other phylogenetic tree comparison frameworks, such as ADView ^15^ and Phylo.io ^23^. The process of comparing two trees with iPhyloC reaches interactive frame rates, even for topologies with a huge number of terminals, which is an enhancement contrasting to other available frameworks.

iPhyloC can handle two large phylogenetic trees. We conducted a scalability test using a MacBook Air (early 2014, 1.7 GHz Dual-Core Intel Core i7 processor, and 8 GB 1600 MHz DDR3 memory). Using the function rtree offered in the R package “ape” ^19,18^, we generated random phylogenetic trees with 80.000 taxa, 90.000 taxa, 100.000 taxa, and 110.000 taxa. iPhyloC was able to handle two trees of up to 100.000 taxa each, but the browser crashed when using two trees with 110.000 taxa and more. The conducted scalability test is not conclusive as it depends on the user’s device specifications, and on the hosting server specifications as well.

The power of iPhyloC comes from our design choice of not forcing the tree to fit the the user’s screen size and from allowing comparison in radial layout, saving more space than the linear layout. Another strength of iPhyloC over other phylogenetic trees comparison frameworks is that the pre-processing of trees is done using fast set based calculations. Further in-depth trees exploration and comparison is carried out in interactive frame rates using JavaScript, which runs in the user’s browser. Additionally, the user can export the visualized trees in Scalable Vector Graphics (SVG) format, which offers very high resolution images in a small file size.

## 8. Conclusion

We tackle the problem of one-to-one tree comparison in the domain of phylogenetic trees analysis through a novel framework named iPhyloC, along with a new comparison technique, the corresponding subtree. Our results were validated by direct comparison with Phylo.io ^23^. Generally, comparison frameworks accept binary and highly overlapping trees only or trees with the same sets of taxa. Here, we consider a usage scenario that demands a different approach: phylogenetic supertrees. Comparing source trees with the inferred supertree is especially hard because, in most cases, the supertree is not fully resolved and might not highly overlap with its source trees. iPhyloC succeeds in such a task.

Further work will extend iPhyloC to deal with one-to-many and general tree comparison problems such as trees with duplicated taxa, especially relevant in gene trees investigations, host-parasite comparisons (a single host with different parasites or a single organism parasitizing different hosts), and historical biogeographical date (with widespread taxa and redundant distributions). Another future direction is to add visual compression technique (e.g. focus+context) to enhance the visual scalability of iPhyloC.

